## Supplementary figures and images for "AnimalGAN: A Generative Adversarial Network Model Alternative to Animal Studies for Clinical Pathology Assessment"

### Figure S1

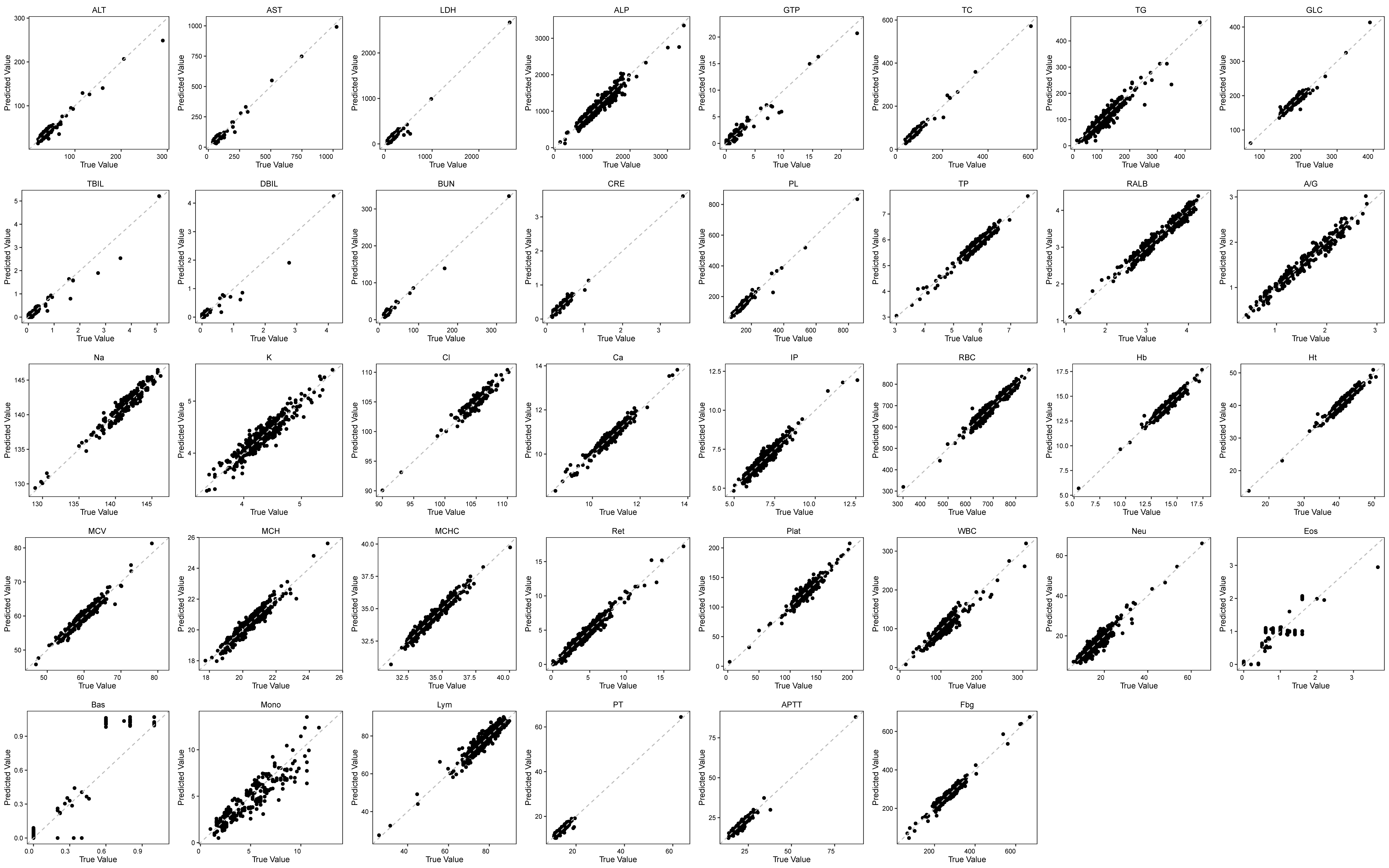

### Figure S2

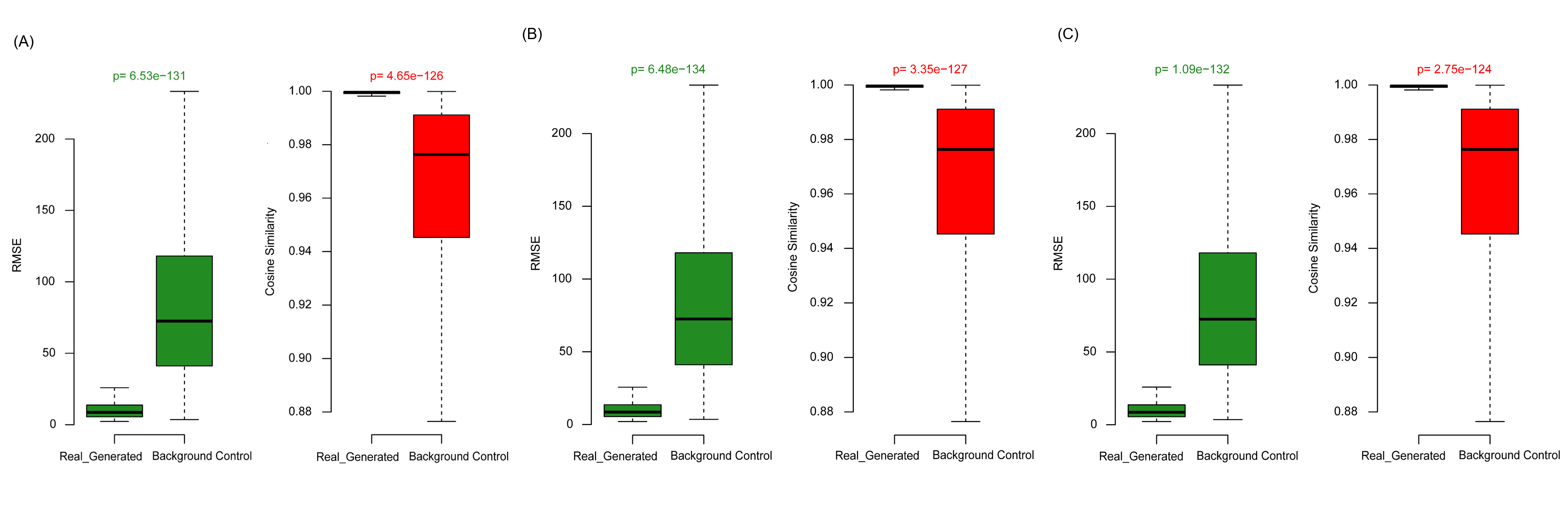

### Figure S3

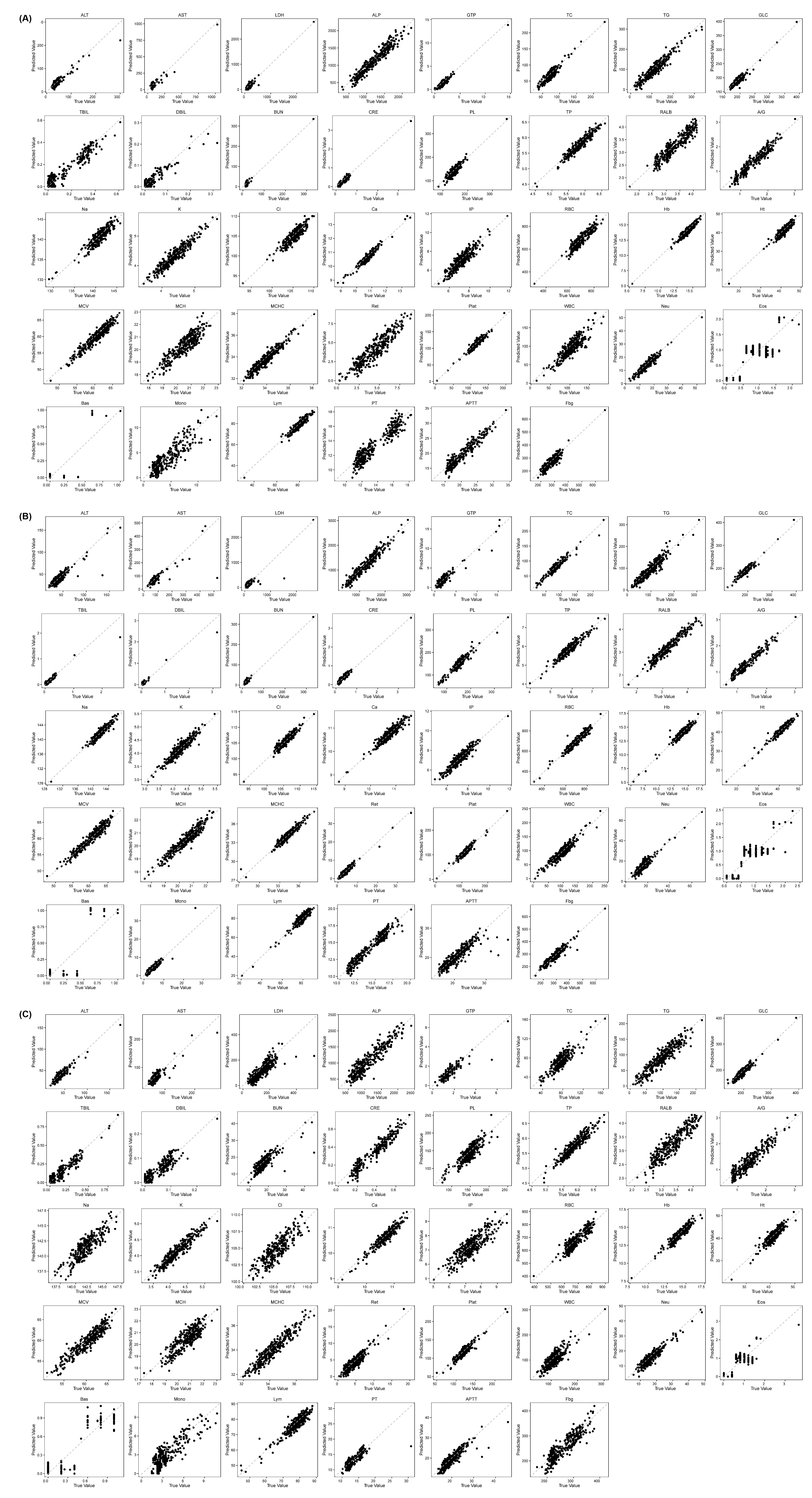

### Figure S4

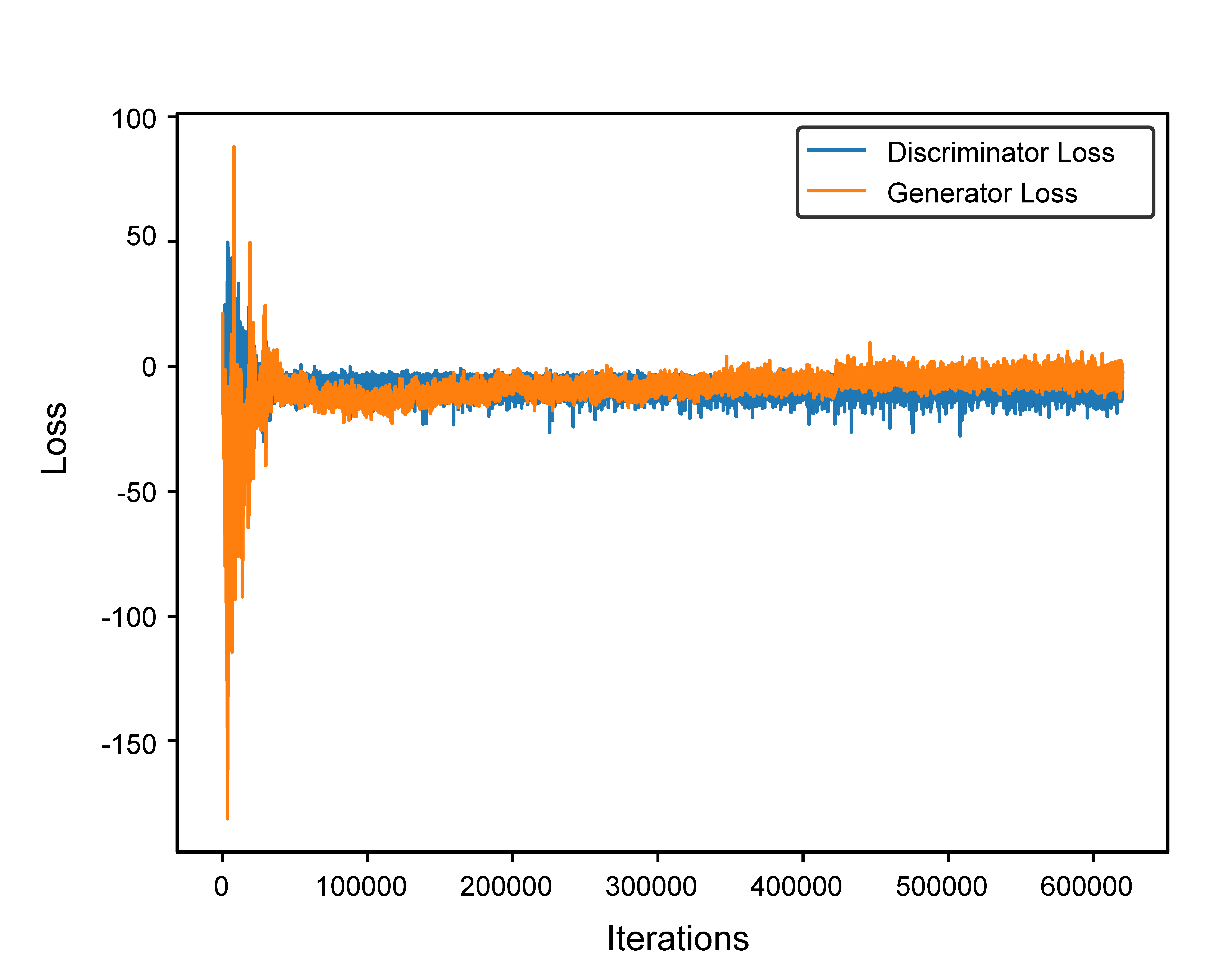

### Figure S5

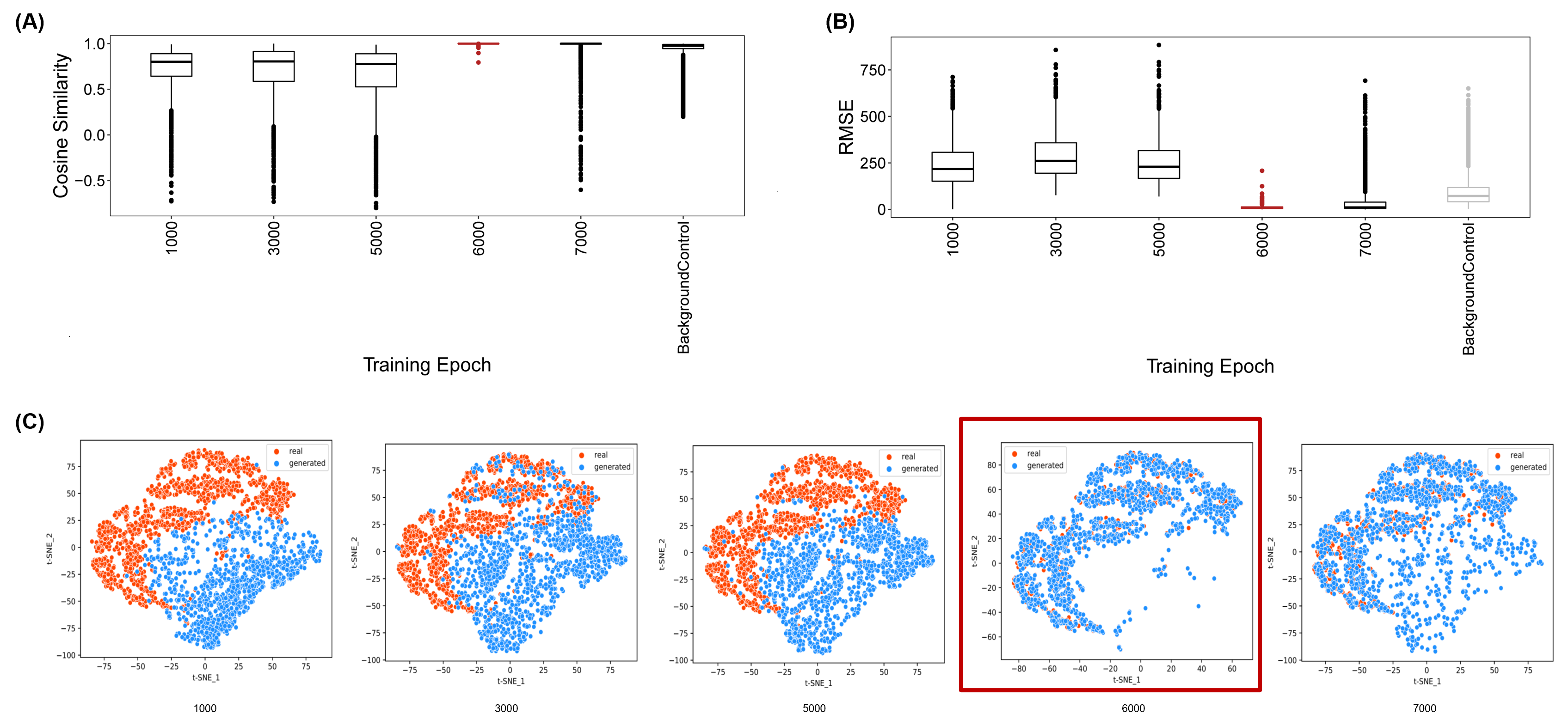

### Figure S6

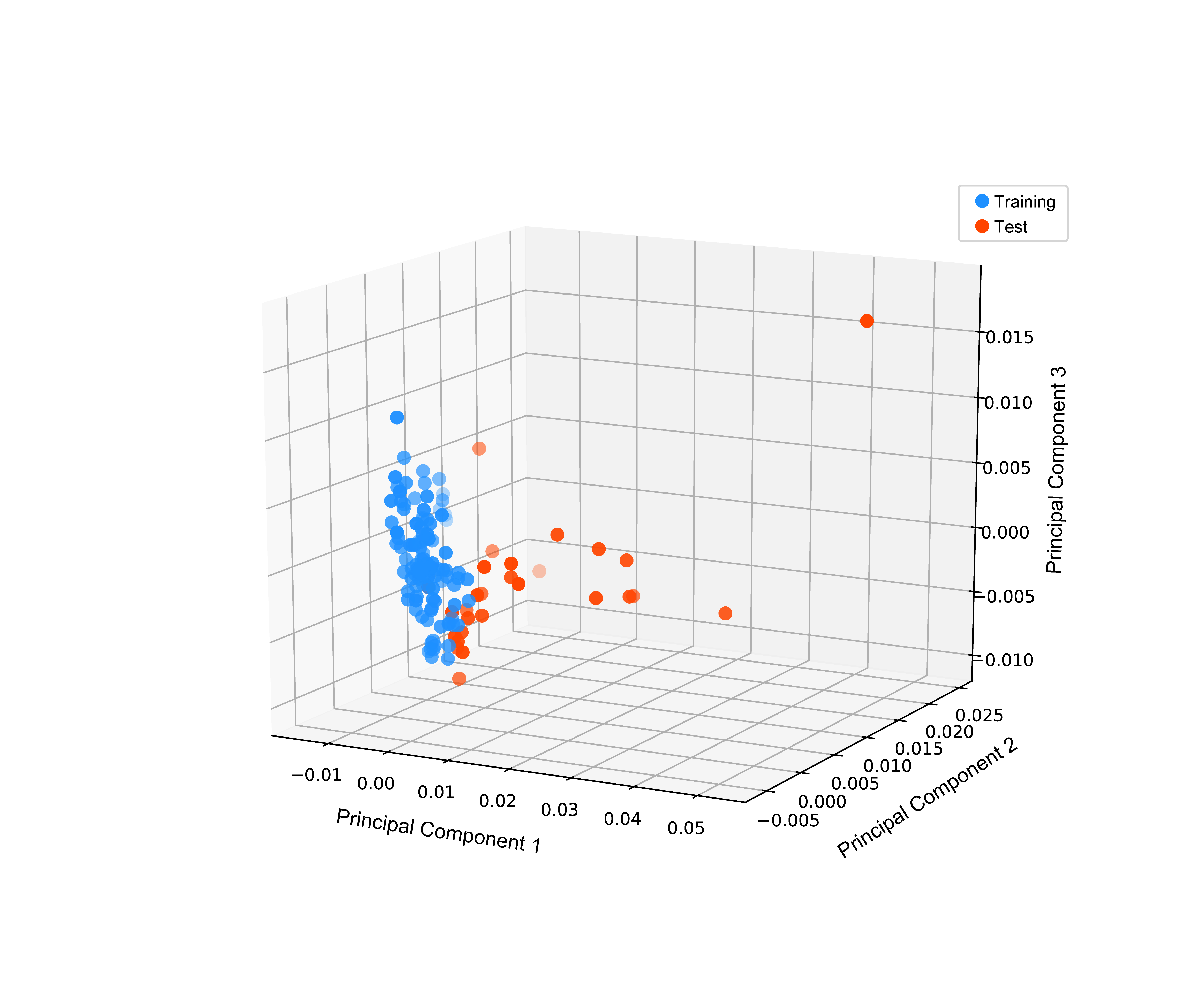

### Figure S7

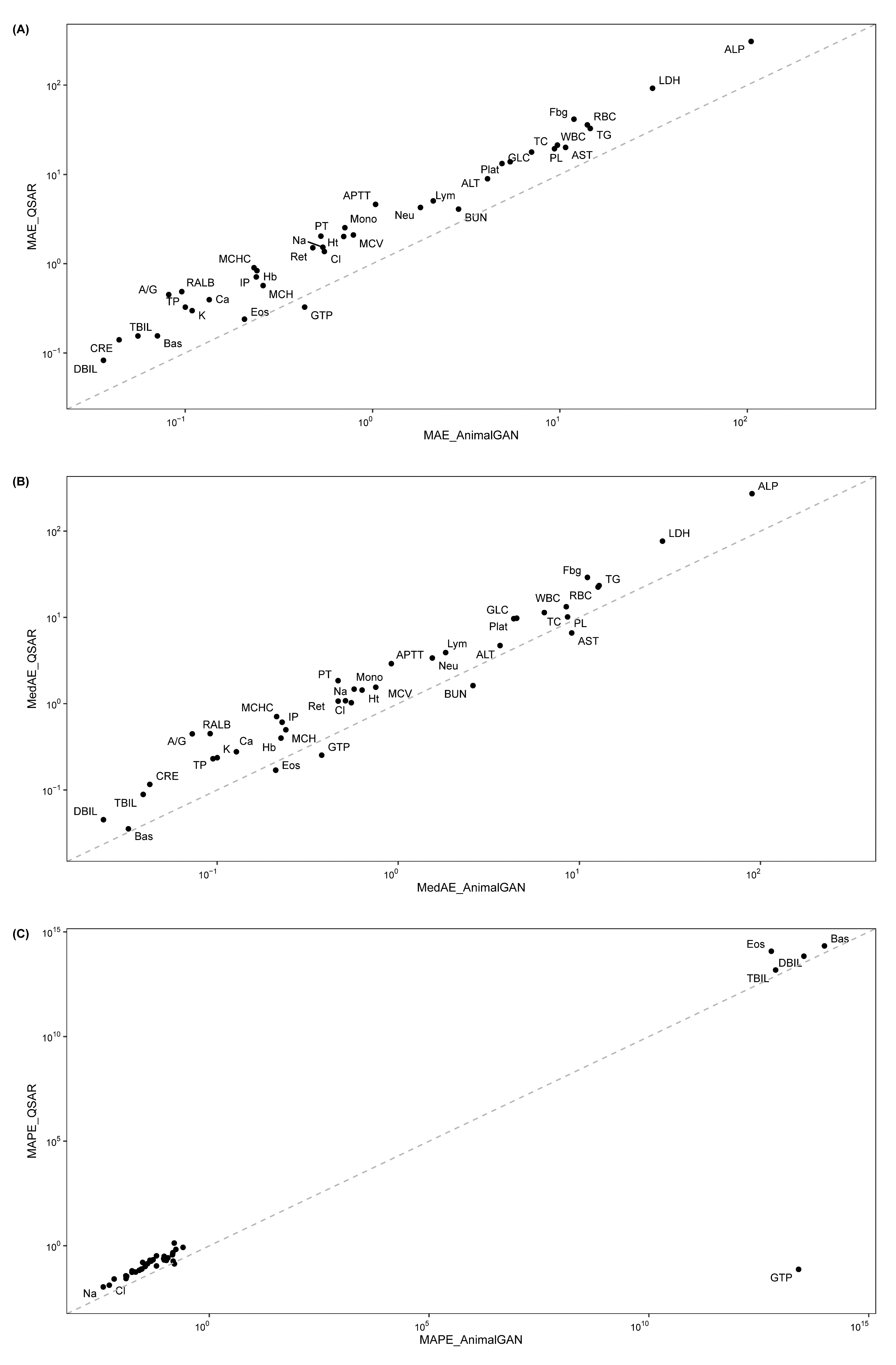
