## Supplementary documents for "AnimalGAN: A Generative Adversarial Network Model Alternative to Animal Studies for Clinical Pathology Assessment"

**Supplementary Information for**

**AnimalGAN Predicts Clinical Pathology and Idiosyncratic Drug-induced Liver Injury**

Xi Chen^1^, Scott Auerbach^2^, Ruth Roberts^3,4^, Zhichao Liu^1,5*^, Weida Tong^1*^

^1^ National Center for Toxicological Research, Food and Drug Administration, Jefferson, AR, USA

^2^ Division of the Translational Toxicology, National Institute of Environmental Health Sciences, Research Triangle Park, North Carolina, USA

^3^ ApconiX Ltd, Alderley Park, Alderley Edge, SK10 4TG, U.K.

^4^ University of Birmingham, Edgbaston, Birmingham, B15 2TT, U.K.

^5^ Currently working at Boehringer Ingelheim

**Disclaimer**

*This manuscript reflects the views of the authors and does not necessarily reflect those of the Food and Drug Administration. Any mention of commercial products is for clarification only and is not intended as approval, endorsement, or recommendation.*

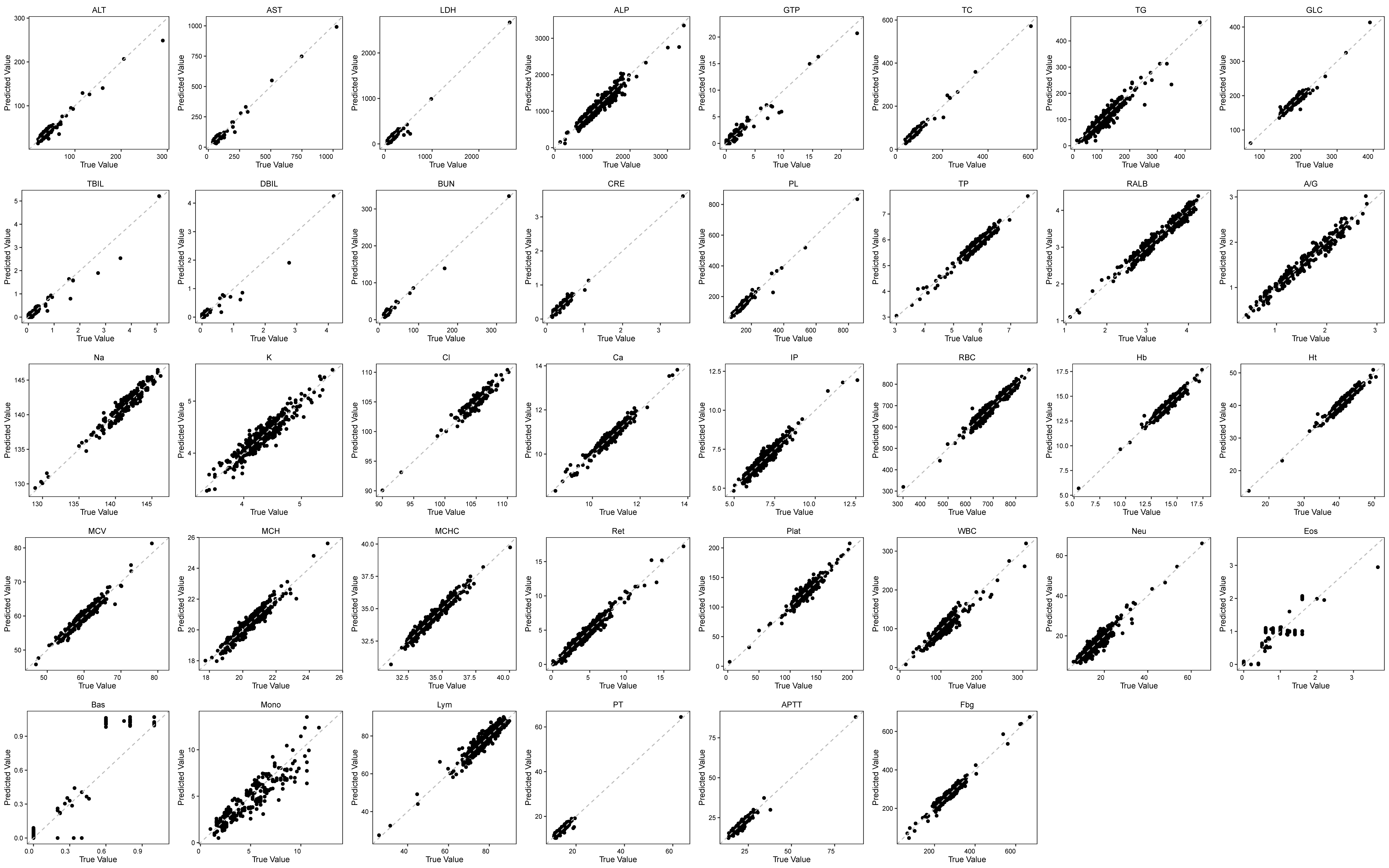

**Supplementary Figure S1. Prediction error plots of AnimalGAN synthetic results against actual laboratory animal testing data for each of the 38 clinical pathology measurements.** Each point represents a treatment condition in the test set. Points on the diagonal depict perfect prediction. Points below the diagonal were underestimated by AnimalGAN (predicted value is lower than the true value), while points above the diagonal were overestimated (predicted value is higher than the true value).

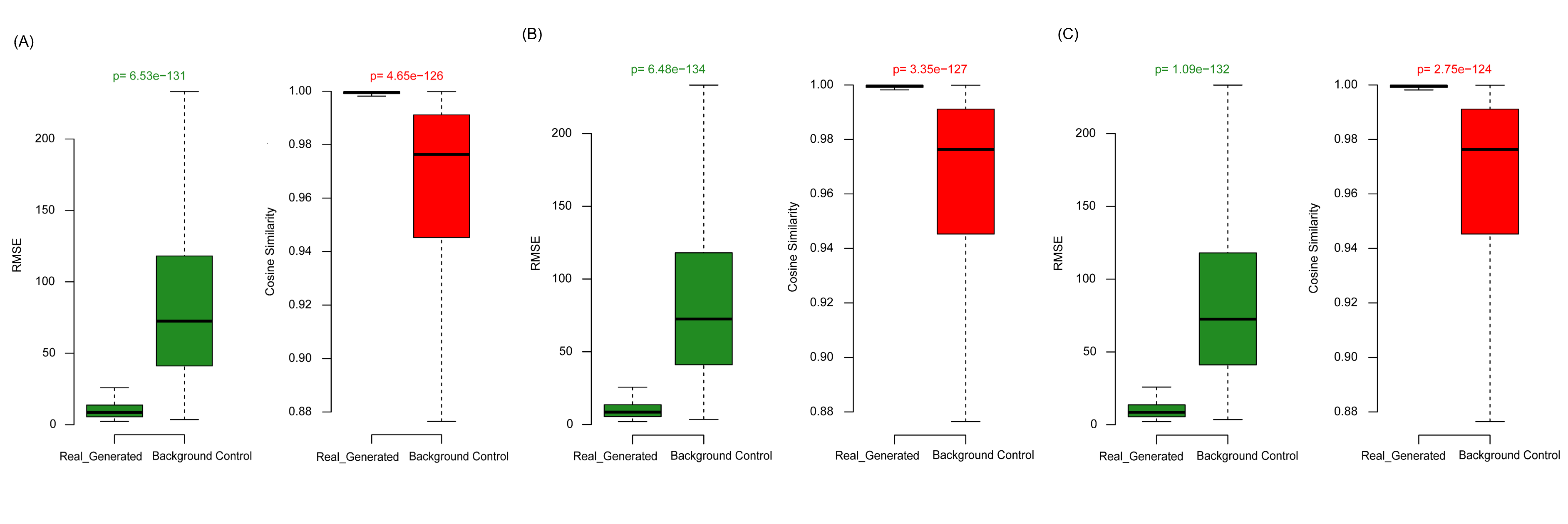

**Supplementary Figure S2. AnimalGAN evaluation in three distinct real-world scenarios.** RMSEs and Cosine Similarities between synthetic data and laboratory animal testing data for treatment conditions in test set comparing with those of background distribution. (A) Keep compounds whose chemical structures were far different from those that were used to develop AnimalGAN model in test set; (B) Keep drugs whose therapeutical classes were not included in the development of AnimalGAN in test set; (C) Keep drugs that were approved by FDA more recently in test set.

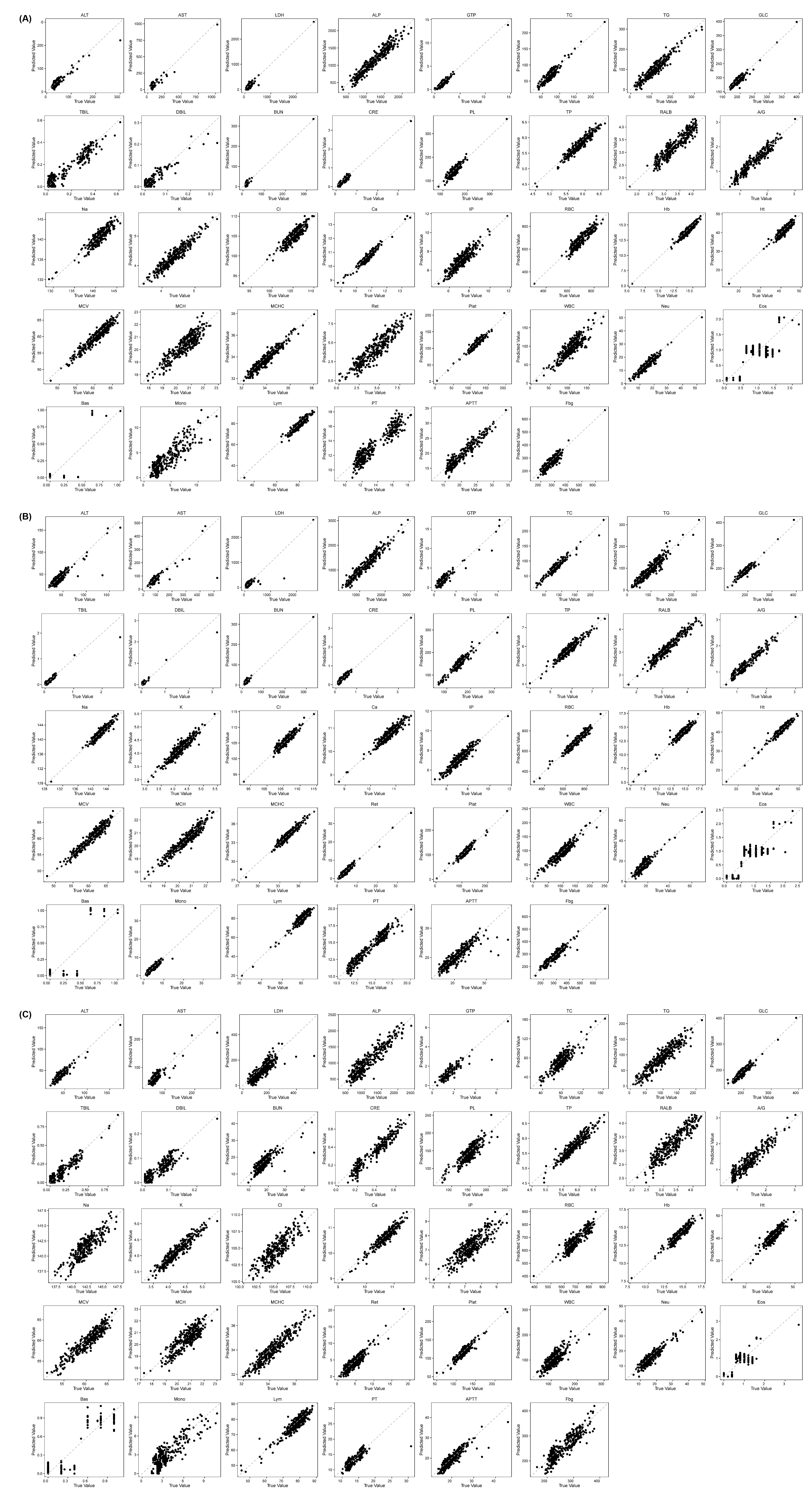

**Supplementary Figure S3. Prediction error plots of synthetic results against actual testing values for each of the 38 clinical pathology measurements for treatment conditions in test set.** (A) Keep compounds whose chemical structures were far different from those that were used to develop AnimalGAN model in test set; (B) Keep drugs whose therapeutical classes were not included in the development of AnimalGAN in test set; (C) Keep drugs that were approved by FDA more recently in test set. Each point represents a treatment condition in test set. Points on the diagonal depict perfect prediction. Points below the diagonal were underestimated (predicted value is lower than the true value), while points above the diagonal were overestimated (predicted value is higher than the true value).

| **Measurement** | **Consistency** | **Toxicity** |
| --- | --- | --- |
| GTP | 0.9608 | Hepatotoxicity |
| LDH | 0.9789 | Hepatotoxicity |
| TBIL | 0.9819 | Hepatotoxicity |
| DBIL | 0.9849 | Hepatotoxicity |
| ALT | 0.9910 | Hepatotoxicity |
| AST | 0.9970 | Hepatotoxicity |
| ALP | 1.0000 | Hepatotoxicity |
| BUN | 0.9789 | Nephrotoxicity |
| K | 0.9880 | Nephrotoxicity |
| CRE | 0.9940 | Nephrotoxicity |
| Na | 1.0000 | Nephrotoxicity |
| Cl | 1.0000 | Nephrotoxicity |
| Ca | 1.0000 | Nephrotoxicity |
| IP | 1.0000 | Nephrotoxicity |
| Eos | 0.9759 |  |
| Mono | 0.9759 |  |
| Neu | 0.9789 |  |
| WBC | 0.9819 |  |
| Bas | 0.9849 |  |
| TG | 0.9880 |  |
| PL | 0.9910 |  |
| Ht | 0.9910 |  |
| Plat | 0.9910 |  |
| TC | 0.9940 |  |
| MCH | 0.9940 |  |
| PT | 0.9940 |  |
| APTT | 0.9940 |  |
| Fbg | 0.9940 |  |
| GLC | 0.9970 |  |
| Ret | 0.9970 |  |
| TP | 1.0000 |  |
| RALB | 1.0000 |  |
| A/G | 1.0000 |  |
| RBC | 1.0000 |  |
| Hb | 1.0000 |  |
| MCV | 1.0000 |  |
| MCHC | 1.0000 |  |
| Lym | 1.0000 |  |

**Supplementary Table S1. The consistency between AnimalGAN results and real animal testing data on toxicity assessment for each of the 38 clinical pathology measurements.** Unpaired t-tests were performed to compare the controls and treatments, and p < 0.05 were considered statistically significant. Calculations were based on the hypothesis that the clinical pathology measurements are normally distributed and that the within-group variances are the same. For those measurements that did not follow the normal distribution, non-parametric tests were used. Then the consistency was calculated based on the comparison of toxicity assessment conclusions between AnimalGAN results and real testing data.

**Supplementary Table S2.** The detailed information on treatment conditions and samples used in this study. Data splitting details in different scenarios are also include in this table.

**Supplementary Table S3. Compounds used in this study.** Compounds details, including SMILES, structural similarity score calculated based on Mordred molecular representations, first level of the WHO Anatomical Therapeutical Chemical (ATC) code, and the initial approval year of each compound.

| Generator *G* | Discriminator *D* |
| --- | --- |
| $\mathcal{z\in}\mathbb{R}^{\mathbf{1828}}\mathcal{\sim N}\left( \boldsymbol{0,I} \right)$; $\mathcal{c}\boldsymbol{\in}\mathbb{R}^{\mathbf{1828}}$ | A sample $\mathcal{x}\in\mathbb{R}^{38}$ with condition $\mathcal{c}\in\mathbb{R}^{1828}$ |
| $\boldsymbol{concat}\left( \mathcal{c, z} \right)\boldsymbol{\in}\mathbb{R}^{\boldsymbol{3656}}$ | $concat\left( \mathcal{c},\mathcal{x} \right)\in\mathbb{R}^{1866}$ |
| Fully connected layer→4096; LeakyReLU | Fully connected layer→2048; LeakyReLU |
| Fully connected layer→2048; LeakyReLU | Fully connected layer→1024; LeakyReLU |
| Fully connected layer→1024; LeakyReLU | Fully connected layer→ 256; LeakyReLU |
| Fully connected layer→256; LeakyReLU | Fully connected layer→64; LeakyReLU |
| Fully connected layer→64; LeakyReLU | Fully connected layer→32; LeakyReLU |
| Fully connected layer→38 | Fully connected layer→1 |

**Supplementary Table S4. Network architectures for the generator and discriminator of AnimalGAN.** The noise $\mathcal{z}$ is set to a vector of the same dimension as the condition $\mathcal{c}$ sampled from a multivariate Gaussian distribution $\mathcal{N}\left( \boldsymbol{0,I} \right)$. The covariance matrix here is the identify matrix of size 1828, denoted by $\boldsymbol{I}$.

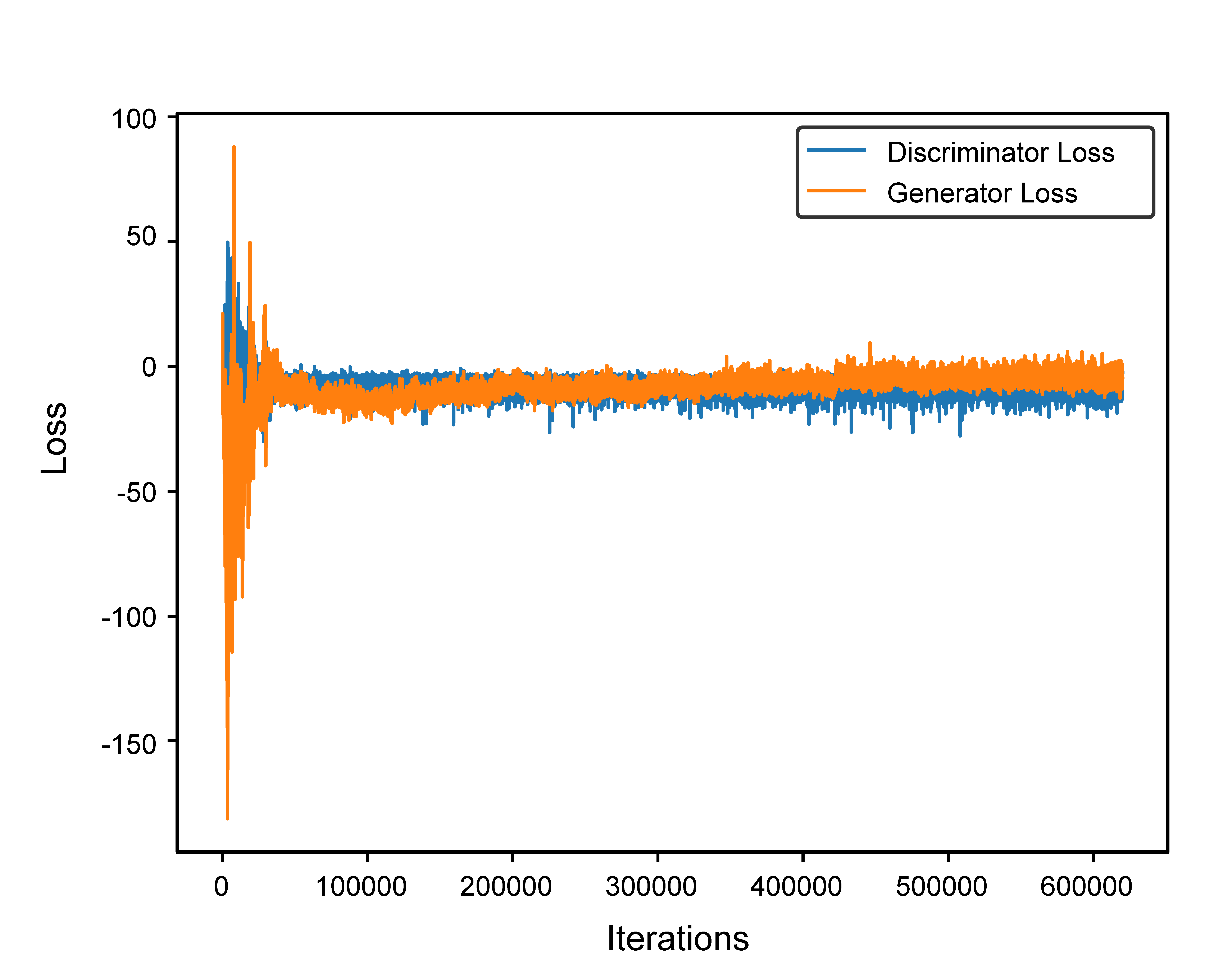

**Supplementary Figure S4. Loss curves of generator and discriminator.**

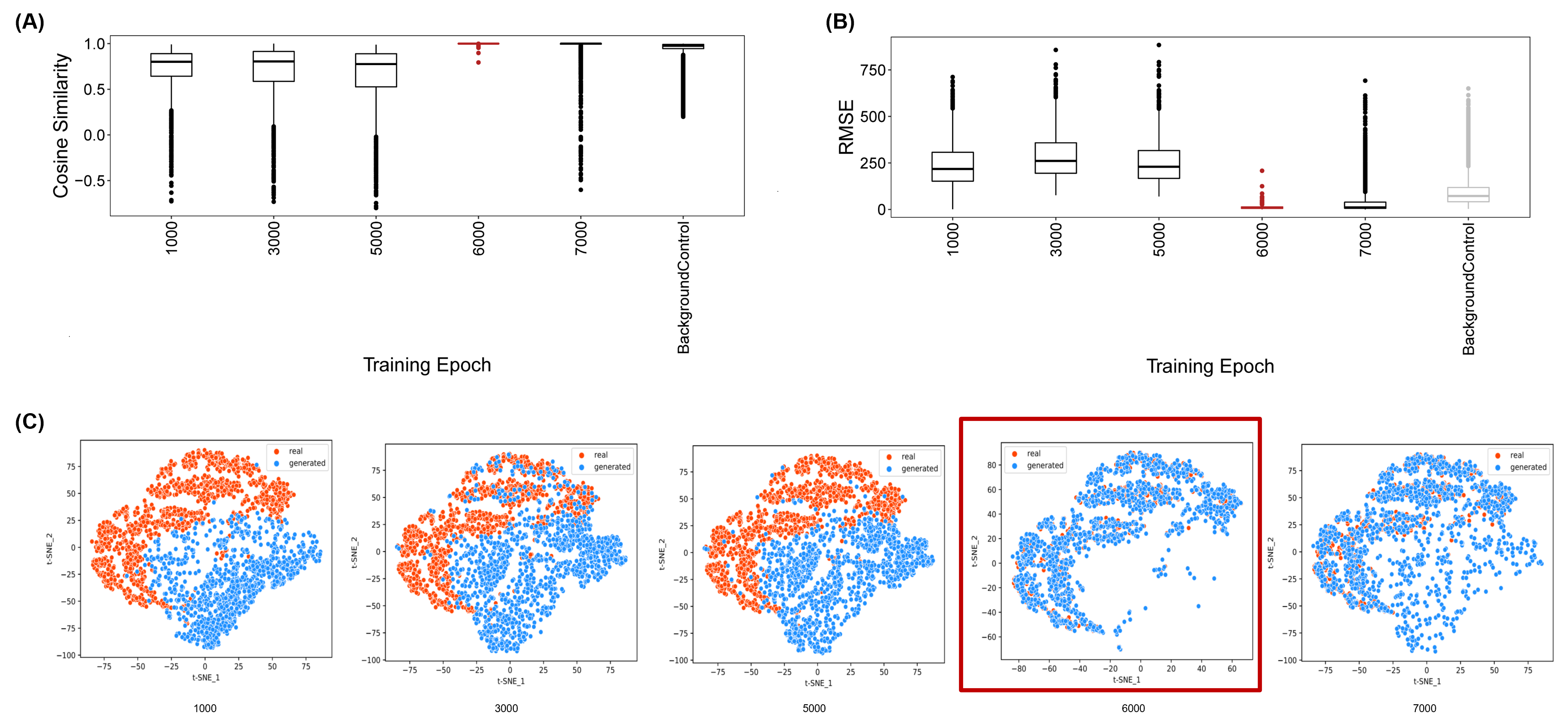

**Supplementary Figure S5. Evolution of model performance during training.** Distributions of (A) cosine similarities and (B) RMSEs between generated data and real animal testing data for all the treatment conditions in training set along with training. (C) t-SNE visualization of generated data and real data for treatment conditions in the training set at different training epochs. Each point depicted one treatment condition.

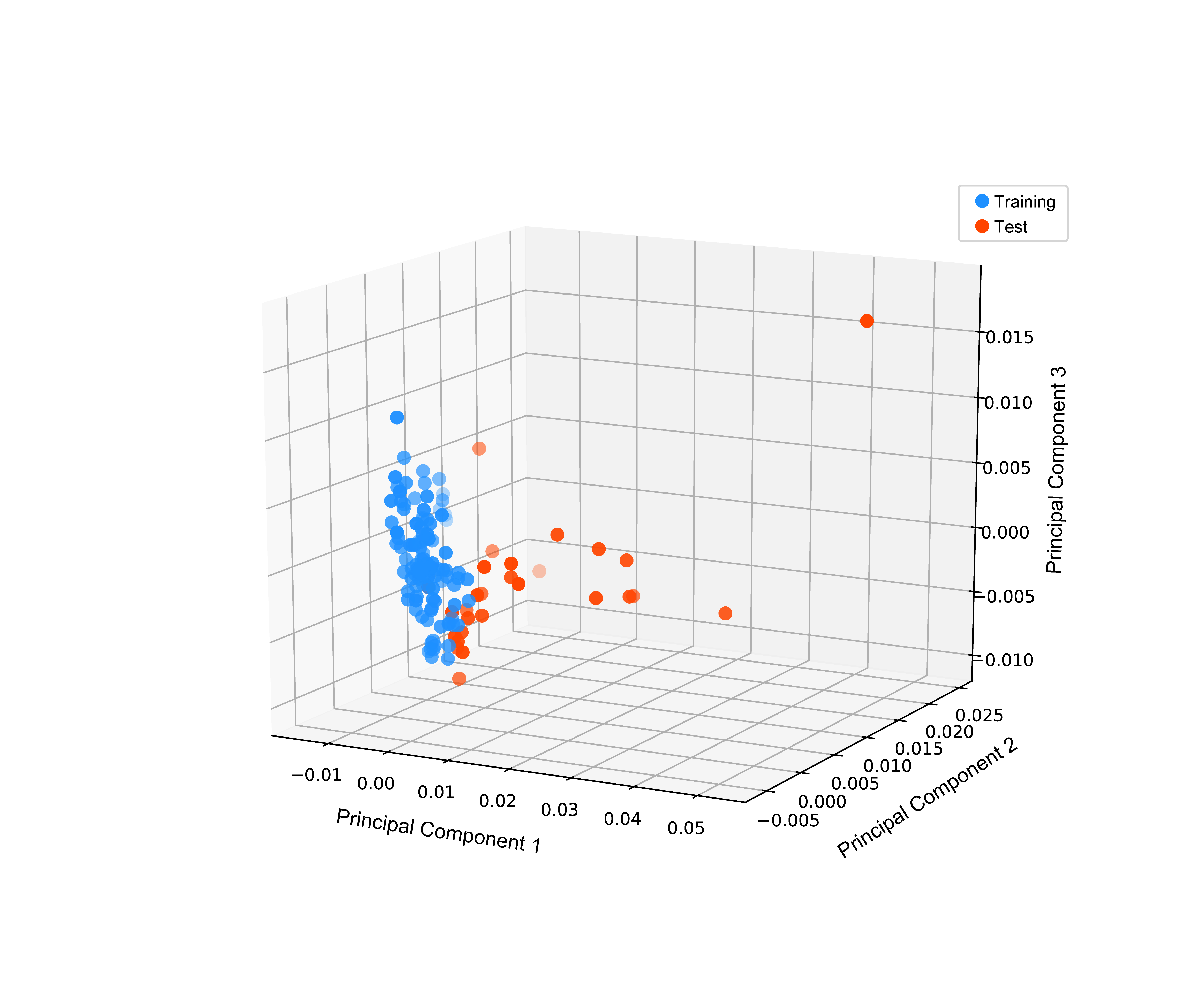
**Supplementary Figure S6. Visualization of structural similarities of all the 138 compounds.** The pairwise structural similarities between any two of the 138 compounds were calculated based on their Mordred molecular representations. Each point depicted one compound.

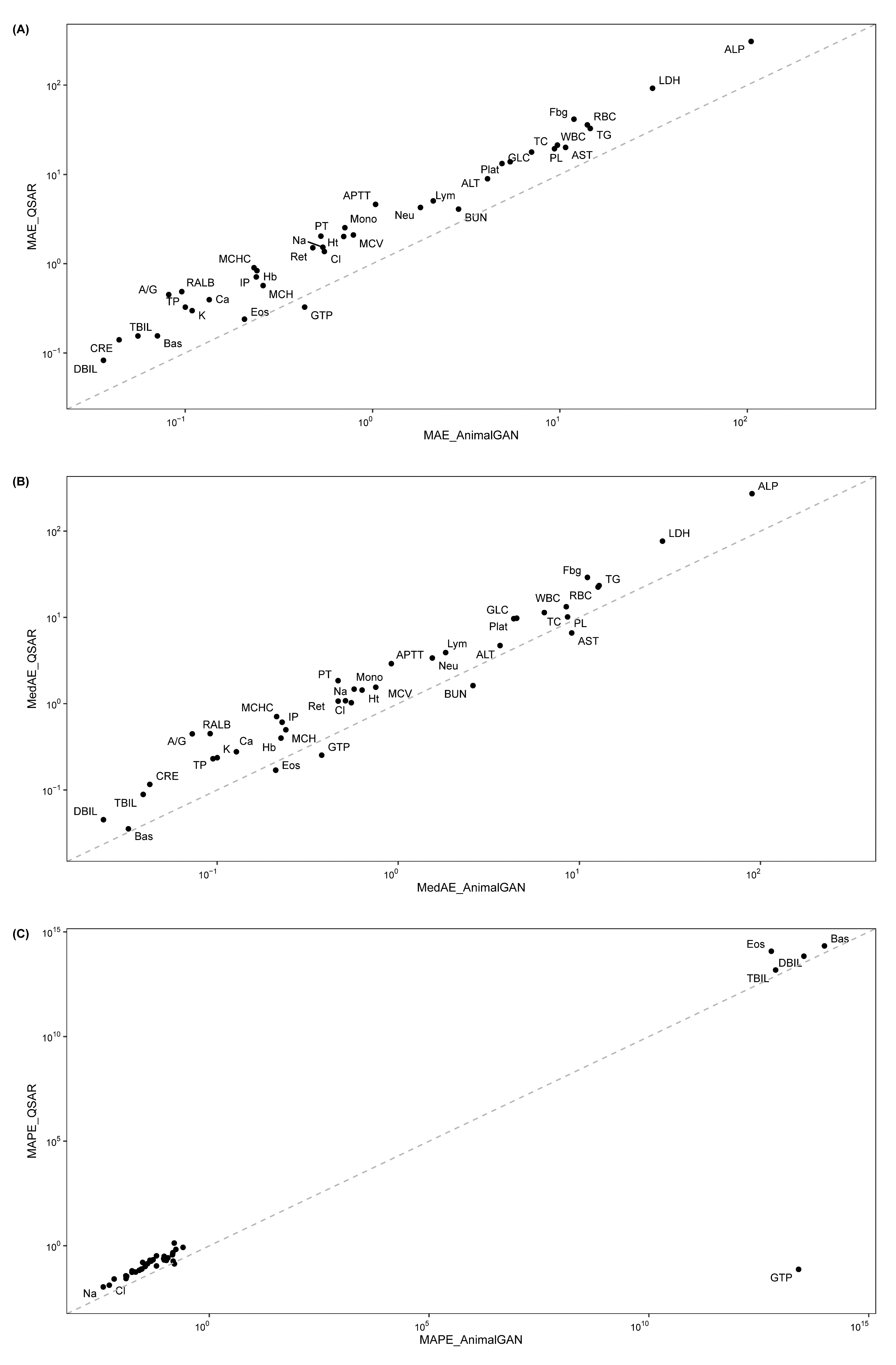

**Supplementary Figure S7. Comparisons of AnimalGAN results with QSAR predictions for all 38 clinical pathology measurements in terms of metrics (A) Mean absolute error (MAE), (B) MedAE (Median Absolute Error), and (C)Mean Absolute Percentage Error (MAPE).** Each point represents a clinical pathology measurement. Points on the diagonal depict the comparable performance between AnimalGAN and QSAR analyses. Points above the diagonal demonstrates AnimalGAN outperforms QSARs. Points below the diagonal demonstrates AnimalGAN underperforms QSARs. All comparisons demonstrate AnimalGAN outperform QSAR analyses for most of the clinical pathology measurements.
